# Gold Nanoparticles for Monitoring of Mesenchymal Stem Cell–Loaded Bioresorbable Polymeric Wraps for Arteriovenous Fistulas

**DOI:** 10.1101/2023.02.01.526611

**Authors:** Allan John R. Barcena, Joy Vanessa D. Perez, Jossana A. Damasco, Marvin R. Bernardino, Erin Marie D. San Valentin, Carleigh Klusman, Benjamin Martin, Andrea Cortes, Gino Canlas, Huckie C. Del Mundo, Francisco M. Heralde, Rony Avritscher, Natalie Fowlkes, Richard R. Bouchard, Jizhong Cheng, Steven Y. Huang, Marites P. Melancon

## Abstract

**Background:** To address high rates of arteriovenous fistula (AVF) failure, a mesenchymal stem cell (MSC)-seeded polymeric perivascular wrap has been developed to reduce neointimal hyperplasia (NIH) and enhance AVF maturation in a rat model. However, the wrap’s radiolucency makes its placement and integrity difficult to monitor.

**Purpose:** In this study, we infused gold nanoparticles (AuNPs) into the polymeric perivascular wrap to improve its radiopacity and tested the effect of infusion on the previously reported beneficial effects of the polymeric wrap on the AVF outflow vein.

**Materials and Methods:** We fabricated a polymeric perivascular wrap made of polycaprolactone (PCL) infused with AuNPs via electrospinning. Sprague-Dawley rat mesenchymal stem cells (MSCs) were seeded on the surface of the wraps. We then compared the effect of five AVF treatments—no perivascular wrap (i.e., control), PCL wrap, PCL+MSC wrap, PCL-Au wrap, and PCL-Au+MSC wrap—on AVF maturation in a Sprague-Dawley rat model of chronic kidney disease (n=3 per group). Statistical significance was defined as p<.05, and one-way analysis of variance was performed using GraphPad Prism software.

**Results:** On micro-CT, AuNP-infused wraps demonstrated significantly higher radiopacity compared to wraps without AuNPs. On ultrasonography, wraps with and without AuNPs equally reduced the wall-to-lumen ratio of the outflow vein, a marker of vascular stenosis. On histomorphometric analysis, wraps with and without AuNPs equally reduced the neointima-to-lumen ratio of the outflow vein, a measure of NIH. On immunofluorescence analysis, representative MSC-seeded wraps demonstrated reduced neointimal staining for markers of smooth muscle cells (α-SMA), inflammatory cells (CD45), and fibroblasts (vimentin) infiltration when compared to control and wraps without MSCs.

**Conclusion:** Gold nanoparticle infusion allows the in vivo monitoring via micro-CT of a mesenchymal stem cell–seeded polymeric wrap over time without compromising the benefits of the wrap on arteriovenous fistula maturation.

**Summary Statement:** Gold nanoparticle infusion enables in vivo monitoring via micro-CT of the placement and integrity over time of mesenchymal stem cell–seeded polymeric wrap supporting arteriovenous fistula maturation.

**Key Results:**

- Gold nanoparticle (AuNP)–infused perivascular wraps demonstrated higher radiopacity on micro-CT compared with wraps without AuNPs after 8 weeks.
- AuNP-infused perivascular wraps equally improved the wall-to-lumen ratio of the outflow vein (a marker of vascular stenosis) when compared with wraps without AuNPs, as seen on US.
- AuNP-infused perivascular wraps equally reduced the neointima-to-lumen ratio of the outflow vein (a measure of neointimal hyperplasia) when compared with wraps without AuNPs, as seen on histomorphometry.

## INTRODUCTION

Approximately 2.3 million individuals are undergoing chronic hemodialysis worldwide, and a majority of them rely on an arteriovenous fistula (AVF) for vascular access (1). In the United States, 62.6% of patients with prevalent end-stage renal disease depend on an AVF for hemodialysis (2). Although an AVF is preferred over alternative forms of vascular access (i.e., intravenous catheter or arteriovenous graft) because of lower rates of infection and maintenance interventions (3), its utility is still limited by high rates of maturation failure (4–6). One of the leading causes of AVF failure is neointimal hyperplasia (NIH), a cellular process brought about by the synergistic action of hemodynamic stress, inflammation, and hypoxia on the vascular wall (7, 8). Although most investigational systemic therapies have failed to demonstrate a clear benefit in improving AVF maturation rates, a wide array of localized perivascular interventions, such as mechanical devices and cell-based therapies, have shown potential in mitigating NIH and AVF failure (9).

In a recent study, the perivascular application of mesenchymal stem cell (MSC)–seeded polymeric scaffolds, in the form of wraps sutured around the AVF, was shown to reduce NIH and enhance AVF maturation (10). However, the polymeric wraps’ radiolucency makes in vivo monitoring challenging. In vivo monitoring is an essential component of medical device surveillance, and in its absence, the safety and efficacy of a medical device are both compromised. Although the wraps are clearly delineated from the surrounding vascular layers on histology, they are not easily delineated on in vivo imaging (i.e., ultrasonography [US] and ^18^F-FDG PET). As a solution, polymeric constructs can be synthesized with metallic nanoparticles to give them radiopacity that will enable the tracking of their placement, integrity, and breakdown over time (11, 12). Among the currently available metallic nanoparticles, gold nanoparticles (AuNPs) have shown excellent tunability, radiopacity, and biocompatibility (13, 14).

In this study, we have synthesized an electrospun polymeric perivascular wrap from AuNP-infused polycaprolactone (PCL), seeded it with MSCs, and compared its effect on AVF maturation against non–AuNP-containing wraps. Additionally, this study aimed to monitor the effect of the polymeric wraps on the AVF outflow vein for a longer period of observation (i.e., 8 weeks) than previously reported, with an initial assessment in as early as 2 weeks. We hypothesized that the incorporation of AuNPs will confer sufficient radiopacity to an MSC-seeded polymeric wrap as seen via micro-CT without affecting the wrap’s beneficial effects on the outflow vein of the AVF.

## MATERIALS AND METHODS

### Synthesis of AuNP-Infused Polymeric Wraps

AuNPs were synthesized following a published protocol by Tian et al. (15). AuNPs and PCL (average M_n_ 80,000, Sigma-Aldrich, St. Louis, MO) were mixed in toluene/MeOH to form a solution with 15% PCL and increasing concentrations of AuNPs—0%, 10%, 20%, and 30%. The resulting solutions were electrospun at 1.0 mL/h and 15 kV using a Spraybase electrospinning system (Avectas, Maynooth, Ireland). Synthesis and characterization of AuNP-infused polymeric wraps, MSC culture and attachment, AVF creation and perivascular wrap application, US, and histological analysis were conducted as described in a prior study (10).

### Characterization of AuNP-Infused Polymeric Wraps

The size and morphology of the AuNPs were determined using a JEM 1010 transmission electron microscope (TEM, JEOL USA, Inc., Peabody, MA). The fiber diameter and pore size of wraps were determined using a Nova NanoSEM scanning electron microscope (SEM, Field Electron and Ion Company, Hillsboro, OR) with an EDAX energy-dispersive spectroscopy system (Ametek, Berwyn, PA). SEM images were captured from random areas to determine the mean fiber diameter and mean pore size of each sample via ImageJ software (National Institutes of Health, Bethesda, MD). Porosity was determined using the calculation method (16) and liquid intrusion method (17). Melting temperature (T_m_) and glass transition temperature (T_g_) were determined using an STA PT 1000 thermogravimetric analyzer (Linseis, Selb, Germany). Ultimate tensile strength was determined using an MTESTQuattro materials testing system (ADMET, Norwood, MA).

### Cell Culture and MSC Attachment

Bone-marrow derived Sprague-Dawley rat mesenchymal stem cells that express red fluorescent protein (RFP-MSCs, P2, Creative Bioarray, Shirley, NY) were cultured in DMEM with 10% FBS and 1% penicillin-streptomycin (Corning, Corning, NY). The wraps were subjected to ethylene oxide gas sterilization and then seeded with 1 x 10^5^ RFP-MSCs. After 48 hours, the samples were imaged using an Eclipse Ti2 fluorescence microscope (Nikon, Melville, NY) to visualize the attachment of RFP-MSCs. The cells for in vivo testing were kept in culture media before surgical application.

### AVF Creation and Perivascular Wrap Application

All rat experiments were approved by the Institutional Animal Care and Use Committee. We created a chronic kidney disease (CKD) model using Sprague-Dawley rats using the two-step 5/6^th^ nephrectomy procedure by Wang et al. (18). After 4 weeks, we then created an AVF model via an end-to-side surgical anastomosis of the external jugular vein with the common carotid artery based on a procedure by Wong et al. (19). The rats were divided into five groups—control AVF (i.e., no wrap), PCL wrap, PCL+MSC wrap, PCL-Au wrap, and PCL-Au+MSC wrap—with 3 rats per group. In the wrap groups, a 1 mm x 5 mm piece of the gas-sterilized cylindrical wrap was placed around the outflow vein and secured using 10-0 nylon sutures. The MSC groups received a wrap seeded with 1 x 10^5^ MSCs.

### X-ray and Micro-CT

Prior to in vivo application, a Skyscan 1276 CMOS micro-CT system (Bruker, Billerica, MA) was used to capture x-ray and micro-CT images of the wraps. The radiopacity of the implanted wraps was then measured using the same machine. The radiopacity of the wraps in Hounsfield units (HU) was determined using CTAn micro-CT image analyzer (Bruker).

### Ultrasonography

A Vevo 2100-LAZR system (FUJIFILM VisualSonics, Toronto, ON, Canada) was used to visualize pulsatile arterial waveforms in the external jugular vein of successfully created AVFs. At weeks 2 and 8, the rats were anesthetized, and a 15-MHz B-mode probe was used to get 6 measurements of the wall thickness and luminal diameter across the outflow vein. Wall-to-lumen ratio was calculated by dividing the average wall thickness by the average luminal diameter.

### Histological Analysis

Rats were euthanized by CO_2_ asphyxiation and cervical dislocation following imaging. The outflow veins were harvested after perfusion for formalin fixation and paraffin embedding. Tissue blocks were cut into 4-μm sections and subjected to hematoxylin and eosin (H&E) staining or immunofluorescence multiplex staining with the following markers: (1) alpha smooth muscle actin (α-SMA), which is highly expressed in smooth muscle cells but also expressed in fibroblasts; (2) CD45, which is highly expressed in leukocytes and a marker of inflammatory infiltration; (3) vimentin, which is highly expressed in fibroblasts but also expressed in vascular smooth muscle cells and lymphocytes; and (4) 4’,6-diamidino-2-phenylindole (DAPI), a nuclear stain. An Aperio LV1 real-time digital pathology system (Leica Biosystems, Buffalo Grove, IL) was used to capture images from the H&E slides, and ImageJ software (NIH) was used for quantification of luminal and neointimal areas. A Leica Versa fluorescent-imaging system (Leica Biosystems) was used to capture images from the immunofluorescent slides, and the Halo image analysis platform (Indica Labs, Albuquerque, NM) was used to perform fluorescent cell counting.

### Statistical Analysis

Quantitative data were presented as means ± standard deviations and analyzed using a one-way analysis of variance with GraphPad Prism software, version 9.0.0 (GraphPad, San Diego, CA). Statistical significance was defined as p<.05.

## RESULTS

### Physicochemical Characteristics of the Polymeric Wraps

Figure 1 shows a representative TEM image of the synthesized AuNPs as well as representative SEM, photographic, x-ray, and micro-CT images of the electrospun polymeric wraps. The synthesized AuNPs had an average diameter of 3.93 ± 0.68 nm. The physicochemical properties of the electrospun polymeric wraps are summarized in Table 1. These properties translated to adequate attachment of MSCs, as seen on fluorescence microscopy (Figure 2). Among the AuNP-infused wraps, PCL-AuNP20 (20% concentration of AuNPs) demonstrated a qualitatively more consistent signal intensity on both x-ray and microCT. Hence, this blend was chosen for the subsequent experiments, termed hereafter as “PCL - Au.”

**Figure 1.**
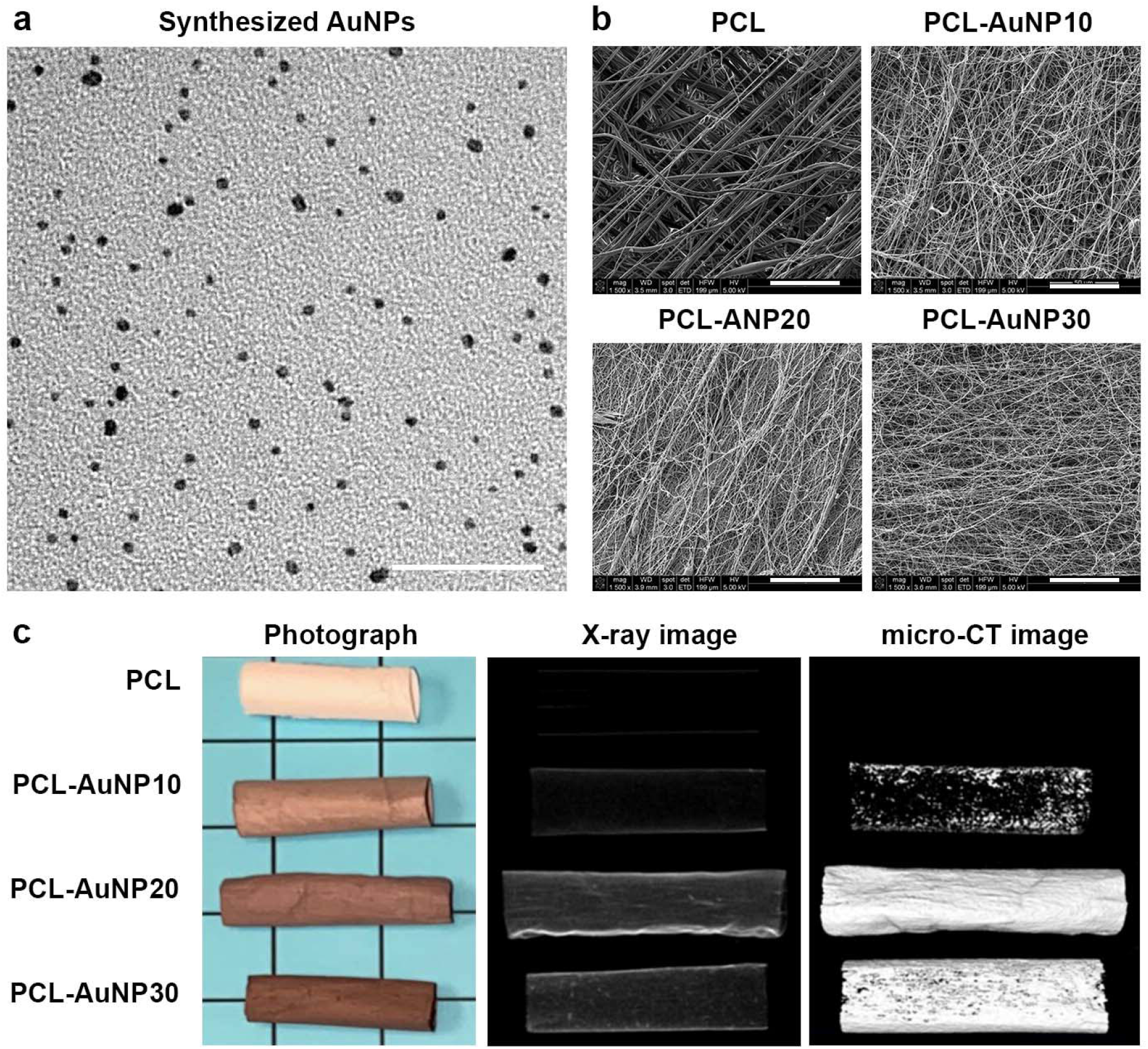
Representative images of the polymeric wraps. (a) Transmission electron microscopy image of synthesized gold nanoparticles (AuNPs) (bar = 50 μm). (b) Representative scanning electron microscopy images of the polymeric wraps (bars = 50 μm). (c) Representative photographic, x-ray, and micro-CT images of the polymeric wraps. Abbreviations: PCL, polycaprolactone.

**Figure 2.**
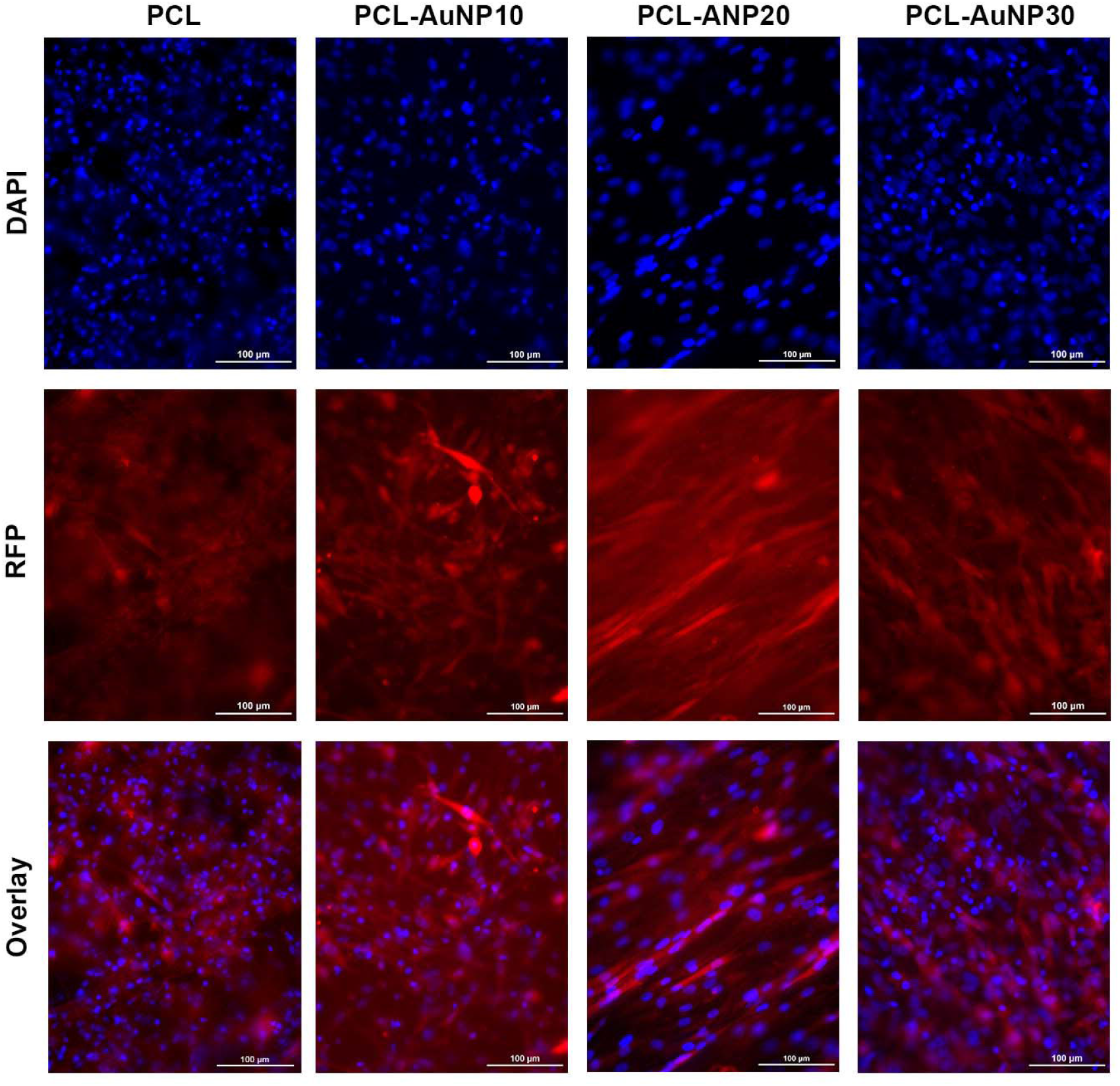
Attachment of Sprague-Dawley rat red fluorescent protein (RFP)–expressing mesenchymal stem cells (MSCs) onto the polymeric wraps. All the polymeric wraps allowed sufficient attachment of Sprague-Dawley rat RFP-MSCs after 48 hours of incubation (bars = 100 μm). Abbreviations: AuNP, gold nanoparticle; DAPI, 4’,6-diamidino-2-phenylindole; PCL, polycaprolactone.

**Table 1.** Physicochemical properties of the electrospun polymeric wraps.

| Wrap type | Diameter ($\mu\text{m}$) | Pore size ($\mu\text{m}$) | Porosity (calculated, %) | Porosity (intrusion, %) | T<sub>g</sub> (°C) | T<sub>m</sub> (°C) | Maximum stress (MPa) |
| --- | --- | --- | --- | --- | --- | --- | --- |
| PCL | 2.07 ± 1.03 | 168.34 ± 177.94 | 93.21 ± 1.23 | 76.75 ± 8.06 | 52.53 | -69.61 | 9.17 ± 0.31 |
| PCL-AuNP10 | 0.53 ± 0.26 | 98.34 ± 76.13 | 53.63 ± 20.62 | 74.31 ± 6.15 | 54.32 | -68.19 | 13.41 ± 0.52 |
| PCL-AuNP20 | 0.25 ± 0.07 | 81.57 ± 86.11 | 65.52 ± 1.13% | 71.13 ± 1.98 | 52.44 | -69.71 | 9.01 ± 0.42 |
| PCL-AuNP30 | 0.27 ± 0.09 | 62.91 ± 53.40 | 64.02 ± 8.34 | 76.03 ± 3.69 | 52.37 | -69.64 | 12.42 ± 1.40 |

Abbreviations: AuNP, gold nanoparticles; PCL, polycaprolactone; T<sub>g</sub>, glass transition
temperature; T<sub>m</sub>, melting temperature.

### AuNP-Infused Wraps Demonstrated Higher Radiopacity on Micro-CT

Figure 3 and Table 2 compare the micro-CT findings for the five groups of rat AVFs 2 and 8 weeks after AVF creation.

**Figure 3.**
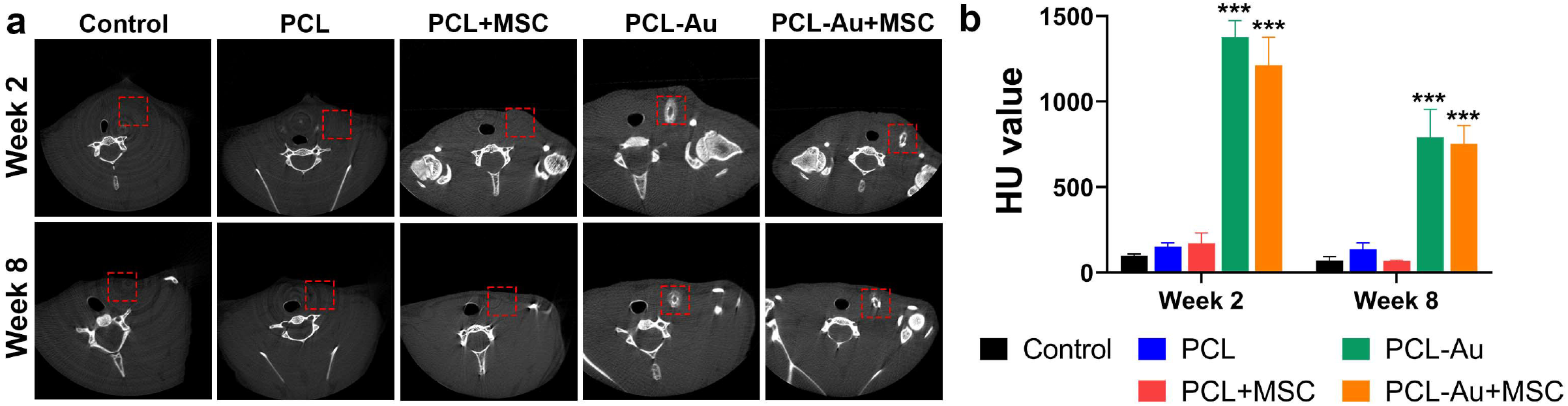
Evaluation of radiopacity via micro-CT. (a) Micro-CT images of the control (i.e., no perivascular wrap), PCL, PCL+MSC, PCL-Au, and PCL-Au+MSC arteriovenous fistulas (AVFs) at 2 and 8 weeks after AVF creation. Red boxes correspond to the position of the wrap. (b) HU values of the five groups of AVFs 2 and 8 weeks after AVF creation; at weeks 2 and 8, rat AVFs wrapped with AuNP-containing polymeric scaffolds (PCL-Au and PCL-Au+MSC) had higher HU values compared to control (p<.001) and non–Au-containing wraps (PCL and PCL+MSC; both p<.001). Abbreviations: ***, p<.001; Au, gold; HU, Hounsfield unit; PCL, polycaprolactone; MSC, mesenchymal stem cell.

**Table 2.** Radiopacity, wall-to-lumen ratio, and neointima-to-lumen ratio of the AVF groups (n=3 per group).

| Wrap type | Radiopacity (HU) |  | Wall-to-lumen ratio |  | Neointima-to-lumen ratio |  |
| --- | --- | --- | --- | --- | --- | --- |
|  | Week 2 | Week 8 | Week 2 | Week 8 | Week 2 | Week 8 |
| Control | 98.07 ± 9.88 | 69.27 ± 23.81 | 0.88 ± 0.44 | 0.95 ± 0.03 | 1.41 ± 0.33 | 4.03 ± 1.9 |
| PCL | 152.0 ± 21.21 | 135.8 ± 37.19 | 0.53 ± 0.05 | 0.31 ± 0.08 | 0.47 ± 0.25 | 0.52 ± 0.23 |
| PCL+MSC | 171.9 ± 60.22 | 67.68 ± 5.61 | 0.5 ± 0.08 | 0.24 ± 0.04 | 1.06 ± 0.59 | 1.06 ± 0.62 |
| PCL-Au | 1375 ± 97.79 | 790.1 ± 164.0 | 0.61 ± 0.09 | 0.33 ± 0.08 | 0.62 ± 0.47 | 1.01 ± 0.64 |
| PCL-Au+MSC | 1212 ± 164.1 | 751.9 ± 107.1 | 0.4 ± 0.09 | 0.23 ± 0.05 | 1.26 ± 0.59 | 0.56 ± 0.37 |

Abbreviations: Au, gold; AVF, arteriovenous fistula; HU, Hounsfield unit; MSC, mesenchymal stem cell; PCL, polycaprolactone

At week 2, higher radiopacity was seen in rat AVFs wrapped with polymeric wraps containing AuNPs (PCL-Au and PCL-Au+MSC) compared to control (p<.001) and non-Au containing wraps (PCL [p<.001] and PCL+MSC [p<.001]). There was no difference among the AVFs without AuNP, i.e., between control and PCL (p=.95), control and PCL+MSC (p=.85), and PCL and PCL+MSC (p>.99). There was also no difference between PCL-Au and PCL-Au+MSC (p=.21) AVFs.

At week 8, the radiopacity of AuNP-containing wraps was retained. PCL-Au and PCL- Au+MSC AVFs still had higher radiopacity compared to control (p<.001), PCL (p<.001), and PCL+MSC (p<.001) AVFs. Similar to week 2, there was no difference between control and PCL (p=.89), control and PCL+MSC (p>.99), or PCL and PCL+MSC (p=.88) AVFs. There was also no difference between PCL-Au and PCL-Au+MSC (p=.98) AVFs. However, there was a reduction in the HU values of the PCL-Au (42.54%, p<.001) and PCL-Au+MSC (37.96%, p=.002) wraps at week 8 when compared to week 2.

### AuNP-Infused and Non-AuNP Wraps Equally Improved the Wall-To-Lumen Ratio of the Outflow Vein

We assessed wall-to-lumen ratio as a marker of vascular stenosis. Figure 4 and Table 2 compare the US findings for the five groups of rat AVFs at 2 and 8 weeks after AVF creation.

**Figure 4.**
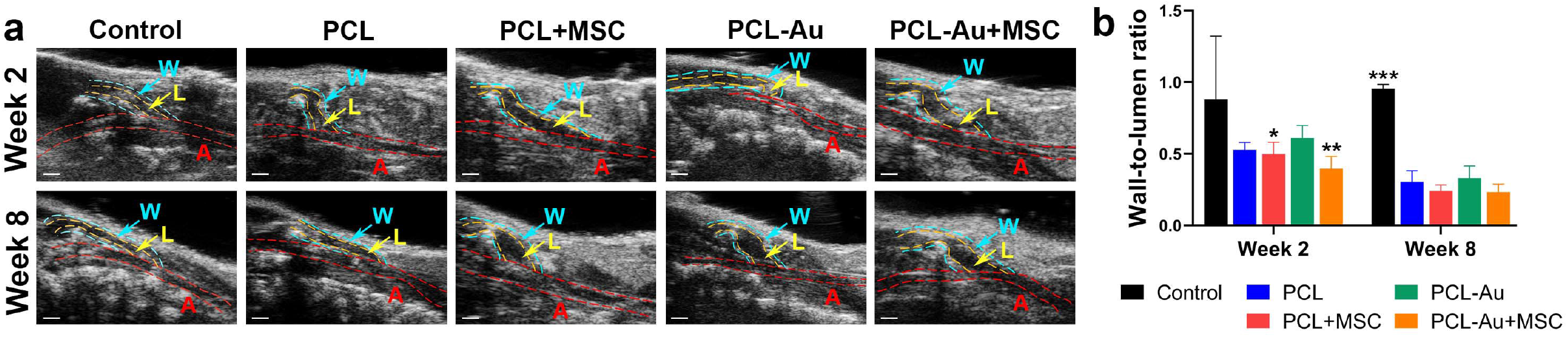
Evaluation of wall-to-lumen ratio via B-mode US. (a) B-mode US images of the control (i.e., no perivascular wrap), PCL, PCL+MSC, PCL-Au, and PCL-Au+MSC arteriovenous fistulas (AVFs) at 2 and 8 weeks after AVF creation (A, artery; L, lumen of outflow vein; W, wall of outflow vein; bars = 1 mm). (b) Wall-to-lumen ratio of the five groups of AVFs 2 and 8 weeks after AVF creation; at week 2, PCL+MSC (p=.045) and PCL-Au+MSC (p=.008) AVFs had a lower wall-to-lumen ratio compared to control, and at week 8, all scaffold groups had a lower wall-to-lumen ratio compared with the control rat AVFs (p<.001). Abbreviations: *, p<.05; **, p<.002; ***, p<.001; Au, gold; PCL, polycaprolactone; MSC, mesenchymal stem cell.

At week 2, the control rat AVFs had a higher wall-to-lumen ratio compared to PCL+MSC (p=.045) and PCL-Au+MSC (p=.008) AVFs. There was no difference between control and PCL (p=.072), control and PCL-Au (p=.24), PCL and PCL+MSC (p>.99), PCL and PCL-Au (p=.96), PCL and PCL-Au+MSC (p=.83), PCL+MSC and PCL-Au (p=.90), PCL+MSC and PCL-Au+MSC (p=.92), or PCL-Au and PCL-Au+MSC (p=.45) AVFs.

At week 8, the control rat AVFs had a higher wall-to-lumen ratio compared to all wrap groups: PCL (p<.001), PCL+MSC (p<.001), PCL-Au (p<.001), and PCL-Au+MSC (p<.001). There was no difference between PCL and PCL+MSC (p=.99), PCL and PCL-Au (p>.99), PCL and PCL-Au+MSC (p=.98), PCL+MSC and PCL-Au (p>.95), PCL+MSC and PCL-Au+MSC (p>.99), or PCL-Au and PCL-Au+MSC (p=.94) AVFs.

### AuNP-Infused and Non-AuNP Wraps Equally Reduced NIH at the Outflow Vein

We assessed neointima-to-lumen ratio to assess the presence of NIH. Figure 5 and Table 2 compare the histomorphometric analysis for the five groups of rat AVFs at 2 and 8 weeks after AVF creation.

**Figure 5.**
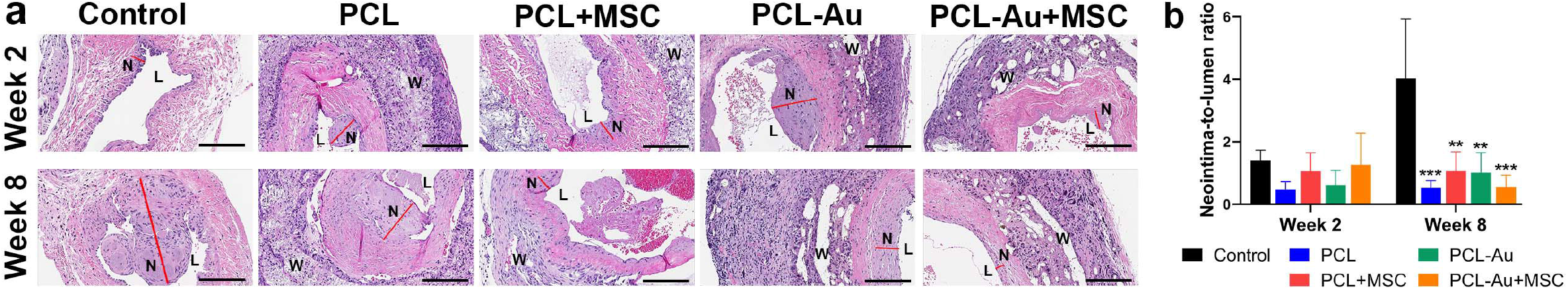
Evaluation of neointima-to-lumen ratio via histomorphometric analysis. (a) Hematoxylin and eosin–stained microscopic images of the control (i.e., no perivascular wrap), PCL, PCL+MSC, PCL-Au, and PCL-Au+MSC arteriovenous fistulas (AVFs) at 2 and 8 weeks after AVF creation (L, lumen; N, neointima; W, wrap; bars = 100 μm). (b) Neointima-to-lumen ratio of the five groups of AVFs 2 and 8 weeks after AVF creation; at week 8, the control rat AVFs had a higher neointima-to-lumen ratio compared to all the other groups. Abbreviations: **, p<.002; ***, p<.001; Au, gold; PCL, polycaprolactone; MSC, mesenchymal stem cell.

**Figure 6.**
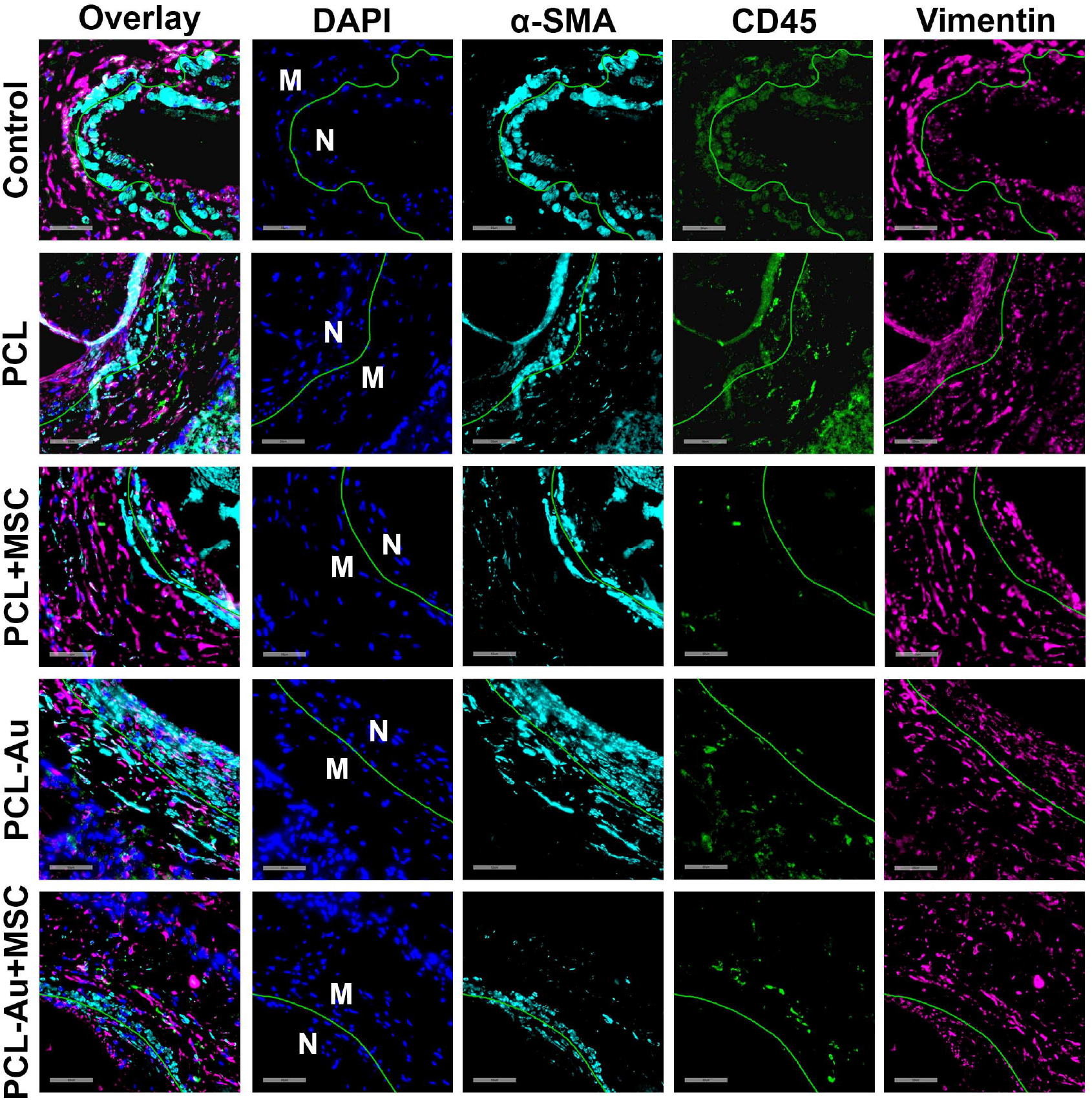
Evaluation of neointimal immunofluorescence staining for α-SMA, CD45, and vimentin. Immunofluorescence microscopic images of the control (i.e., no perivascular wrap), PCL, PCL+MSC, PCL-Au, and PCL-Au+MSC arteriovenous fistulas (AVFs) at 8 weeks after AVF creation, showing the different cell populations in the neointimal (N) and medial (M) vascular layers (bars = 50 μm). Abbreviations: Au, gold; α-SMA, alpha smooth muscle actin; CD, cluster of differentiation; DAPI, 4’,6-diamidino-2-phenylindole; MSC, mesenchymal stem cell; PCL, polycaprolactone.

At week 2, there was no difference in the neointima-to-lumen ratios when comparing any groups: control and PCL (p=.71), control and PCL+MSC (p=.93), control and PCL-Au (p=.86), control and PCL-Au+MSC (p>.99), PCL and PCL+MSC (p>.99), PCL and PCL-Au (p>.99), PCL and PCL-Au+MSC (p=.86), PCL+MSC and PCL-Au (p>.99), PCL+MSC and PCL-Au+MSC (p=.98), PCL-Au and PCL-Au+MSC (p=.95) AVFs.

At week 8, the control rat AVFs had a higher neointima-to-lumen ratio compared to all the other groups: PCL (p=.001), PCL+MSC (p=.002), PCL-Au (p=.002), and PCL-Au+MSC (p<.001). There was no difference between PCL and PCL+MSC (p=.93), PCL and PCL-Au (p=.95), PCL and PCL-Au+MSC (p>.99), PCL+MSC and PCL-Au (p>.99), PCL+MSC and PCL- Au+MSC (p=.94), or PCL-Au and PCL-Au+MSC (p=.96) AVFs.

### MSC-Seeded Wraps Reduced Neointimal α-SMA, CD45, and Vimentin Staining

We assessed several markers that represent neointimal infiltration with smooth muscle cells, inflammatory cells, and fibroblasts (i.e., α-SMA, CD45, and vimentin, respectively). Table 3 compares the immunofluorescence analysis for representative slide from each of the five groups of rat AVFs at 8 weeks after AVF creation. When compared to control, PCL and PCL-Au wraps reduced the neointima-to-media ratio of total cells (i.e., DAPI) as well as CD45- and vimentin-positive cells but increased the neointima-to-media ratio of α-SMA–positive cells. In contrast, PCL+MSC and PCL-Au+MSC wraps reduced the neointima-to-media ratio of α-SMA– positive cells, and these MSC-seeded wraps further decreased the neointima-to-media ratio of total cells as well as CD45- and vimentin-positive cells when compared to their non-MSC counterparts.

**Table 3.** Neointima-to-media ratio of immunofluorescence-stained cells on representative AVFs.

| Wrap type | α-SMA | CD45 | Vimentin | DAPI |
| --- | --- | --- | --- | --- |
| Control | 1.0 | 0.76 | 0.74 | 0.79 |
| PCL | 1.21 | 0.30 | 0.71 | 0.69 |
| PCL+MSC | 0.98 | 0.17 | 0.09 | 0.64 |
| PCL-Au | 1.74 | 0.08 | 0.36 | 0.66 |
| PCL-Au+MSC | 0.92 | 0.07 | 0.22 | 0.40 |

Abbreviations: α-SMA, alpha smooth muscle actin; Au, gold; AVF, arteriovenous fistula; CD, cluster of differentiation; DAPI, 4',6-diamidino-2-phenylindole; MSC, mesenchymal stem cell; PCL, polycaprolactone.

## DISCUSSION

Prior studies have shown that a mesenchymal stem cell (MSC)–seeded polymeric wrap is capable of reducing neointimal hyperplasia (NIH) and enhancing arteriovenous fistula (AVF) maturation (10). However, the radiolucent nature of the polymeric wrap makes in vivo monitoring of its deployment and structural integrity difficult. To address this limitation, we incorporated AuNPs in the synthesis of the polycaprolactone (PCL) wraps, which turned the wraps more radiopaque than their non-AuNP counterparts and consequently clearly observable on micro-CT at up to 8 weeks. This property is useful in AVF surveillance since it facilitates quick localization of the AVF for concurrent study using other imaging modalities, such as US or ^18^F-FDG PET. The integration of the AuNPs within the architecture of the wraps facilitates the confirmation of device placement after surgery, and the reduction in detected intensity from the wrap may serve as a marker of its degradation over time. By week 8, the HU values of the PCL- Au and PCL-Au+MSC wraps had dropped by 42.54% and 37.96%, respectively, compared with week 2. Nonetheless, the intensity level of PCL-Au and PCL-Au+MSC wraps remained similar (p=.98), which suggests that the addition of MSCs did not affect wrap degradation over time.

We found that PCL+MSC and PCL-Au+MSC AVFs demonstrated an improved wall-to-lumen ratio, a marker of vascular stenosis, compared with AVFs without MSCs in as early as 2 weeks after AVF creation. This is consistent with the results of a prior study, wherein the addition of MSCs to polymeric wrap reduced the AVF wall-to-lumen ratio when compared to polymeric wrap alone at 4 weeks after AVF creation (10). These results highlight the unique role of MSCs in improving AVF maturation. The similarity between the wall-to-lumen ratios of PCL+MSC and PCL-Au+MSC AVFs (p=.92) also suggests that the incorporation of AuNPs does not affect the activity of MSCs in reducing AVF wall-to-lumen ratio. By week 8, all treatment groups demonstrated an improved wall-to-lumen ratio, but MSCs no longer conferred an advantage. Nonetheless, the similarity between the wall-to-lumen ratios of AVFs with and without MSCs still supports the idea that the incorporation of AuNPs does not interfere with the activity of the wraps in reducing an AVF’s wall-to-lumen ratio.

We also found that all treatment groups demonstrated improved neointima-to-lumen ratios, a measure of NIH, versus control rats by 8 weeks after AVF creation. Our findings show that the reduction in the neointima-to-lumen ratio by PCL and PCL+MSC treatment versus control seen at 4 weeks after AVF creation in a previous study (10) can be maintained for up to 8 weeks. Moreover, the incorporation of AuNPs did not affect the ability of the wraps to reduce the AVF neointima-to-lumen ratio. Although it appears that MSC-containing wraps no longer provided an added benefit by week 8, there could be changes in the cellular and extracellular composition of the vascular wall attributable to MSCs that were not reflected on the results of in vivo imaging. Interestingly, we found that representative MSC-seeded wraps performed better than control and non-MSC counterparts in reducing the neointima-to-media ratio of total cells as well as α-SMA–, CD45-, and vimentin-positive cells on immunofluorescence analysis. Moreover, we observed an increase in the neointima-to-media ratio of α-SMA–positive cells with PCL and PCL-Au AVFs, which indicates inward migration of smooth muscle cells within the neointima after longer periods of observation. Taken together, these findings support the hypothesis that the addition of MSCs may promote better outward vascular remodeling compared to a vascular wrap that serves purely as a physical support.

The scope of this study was limited to assessment of the signal intensity of the wraps on micro-CT and the effect of the wraps on the AVF wall-to-lumen ratio as seen on US and the neointima-to-lumen ratio as seen on histology. Further studies are recommended to explore the relationship between the rate of wrap degradation and its impact on the ability of the wrap to provide attachment to seeded MSCs as well as mechanical support to the maturing AVF. Moreover, there is a need for additional research on the cellular and extracellular composition of the vascular wall in order to determine the fate of the seeded MSCs, understand the effect of MSCs on the different cells of the outflow vein, and identify potential sources of differences between in vivo and histological observations in this study.

Our results indicate that AuNP infusion permits in vivo monitoring via micro-CT of the placement and integrity over time of an MSC–seeded polymeric wrap while preserving the positive effects of the wrap on AVF maturation. Our findings also confirm the role of MSCs in promoting outward vascular remodeling in the setting of AVF maturation. In conclusion, this study provides a basis for the development and clinical use of an MSC-loaded AuNP-infused perivascular wrap to improve AVF maturation and patency.

## ACKNOWLEDGMENTS

The authors would like to acknowledge Sunita C. Patterson in MD Anderson’s Research Medical Library for editing the manuscript, Dr. James Gu at the Electron Microscopy Core at Houston Methodist Research Institute for assisting with the conduct of scanning electron microscopy, Dunn Lab personnel (i.e., Amanda McWatters, Malea L. Williams, and Steve D. Parrish) for assisting with animal experiments, and Small Animal Imaging Facility personnel for assisting with animal imaging.

## Abbreviations

AuNP: gold nanoparticle;
AVF: arteriovenous fistula;
MSC: mesenchymal stem cell;
NIH: neointimal hyperplasia;
PCL: polycaprolactone

